# Integrin α6β4 signaling switches DNA repair from homologous recombination to non-homologous end-joining pathway to sensitize breast cancer cells to cisplatin

**DOI:** 10.1101/785873

**Authors:** Min Chen, Brock Marrs, Lei Qi, Teresa Knifley, Stuart G. Jarrett, Heidi L. Weiss, Rachel L. Stewart, John A. D’Orazio, Kathleen L. O’Connor

## Abstract

Integrin α6β4 is highly expressed in triple negative breast cancer (TNBC) and drives aggressiveness by stimulating proliferation, angiogenesis, cell migration, invasion and metastasis. Signaling from this integrin stimulates DNA repair and apoptosis resistance, suggesting that it could contribute to therapeutic resistance. Upon testing this hypothesis, we found that integrin α6β4 signaling promoted a three-fold greater sensitivity to cisplatin but exhibited no difference in response to other chemotherapies tested. Mechanistic investigations revealed that integrin α6β4 stimulated quicker and higher amplitude of activation of ATM, Chk2, p53, and 53BP1, which required the integrin β4 signaling domain. Genetic manipulation of gene expression demonstrated that mutant p53 cooperated with integrin α6β4 for cisplatin sensitivity and was necessary for downstream phosphorylation of 53BP1 and enhanced ATM activation. Additionally, we discovered that integrin α6β4 preferentially activated DNA-PKc in response to cisplatin, which led to formation of DNA-PKc-p53 complexes and 53BP1 activation. As a result, integrin α6β4 shifted double strand break repair from homologous recombination (HR) to non-homologous end joining (NHEJ). In summary, we discovered a novel function of integrin α6β4 in switching DSB repair from HR to NHEJ that results in cisplatin sensitivity in TNBC.

## Introduction

Due to a shortage of effective targeted therapy (1,2) and aggressive clinical course (3), triple negative breast cancer (TNBC) remains the most lethal breast cancer subtype. Therefore, novel strategies are actively being sought to treat this aggressive breast cancer subtype. The FDA recently approved PARP-1 inhibitor for the treatment of TNBC patients with BRCA mutation (4) and immunotherapy for the treatment of patients with PD-L1-positive, unresected locally advanced or metastatic TNBC, thus creating the first targeted therapies for TNBC. For most TNBC patients, however, therapies that caused DNA damage remain the standard-of–care. These therapies include ionizing radiation (IR), topoisomerase inhibitors (doxorubicin), alkylating agents (cyclophosphamide), nucleoside analogs (capecitabine, gemcitabine) and platinum agents (cisplatin and carboplatin), with platinum agents used most often in metastatic settings (5,6). How the tumor microenvironment contributes to response to these therapies remains poorly understood. Here, we find that integrin α6β4 promotes DNA repair response and in doing so alters cellular responses to DNA damage-inducing chemotherapeutics.

Integrin α6β4 is a laminin receptor that is highly expressed in TNBC, more so than in hormone-positive or HER2-amplified breast cancers (7). Furthermore, it is highly expressed prominently in the basal-like breast cancer subtype (7), which represents about 80% of TNBCs. Integrin α6β4 coordinates and amplifies signals from the microenvironment to drive the most aggressive traits of TNBC by stimulating proliferation, angiogenesis, apoptosis resistance, migration, invasion (8-14) and metastasis (15,16). Early investigation on how integrin α6β4 contributes to carcinoma progression linked its signaling to p53. In a wild–type (wt) p53 background, integrin α6β4 stimulates p53 leading to p21 upregulation, cleavage of Akt and subsequent apoptosis (17-19). In a mutant or null p53 background, however, integrin α6β4 enhances cell survival through stimulation of the PI3K/Akt pathway (17). These aspects of integrin α6β4 signaling along with other aggressive properties led to the concept that integrin α6β4 would alter therapeutic response (20). However, the impact of integrin α6β4 on therapeutic outcome, its signaling to p53, and how it can cooperate with mutant p53 gain-of-function properties, have gone largely unexplored.

Our recent work demonstrated that integrin α6β4 signaling epigenetically regulates the expression of pro-invasive genes by stimulating the base excision repair (BER) pathway leading to promoter DNA demethylation and can enhance UV-induced nucleotide excision repair (NER) (21). In this study, we find that the ability of integrin α6β4 to stimulate NER extends to TNBC cells. Interestingly, integrin α6β4 signaling does not contribute to therapeutic resistance, but rather to specific sensitivity to cisplatin. We trace this effect to the ability of integrin α6β4 to signal through mutant p53, amplify ATM and DNA-PKc activity, increase 53BP1 phosphorylation, and switch double strand break repair from HR to NHEJ. Together, these studies place the integrin α6β4 signaling cascade as an important regulator of genomic stability and an important therapeutic determinant in TNBC.

## Materials and Methods

### Cell lines and drug treatments

BT549 and MDA-MB-231 cells were obtained from American Type Culture Collection (ATCC). MDA-MB-435 cells (clones 6D7 and 3A7) were described in (9). BT549 cells were cultured in RPMI 1640 containing 50 µg/ml insulin (Sigma Aldrich, St. Louis, MO). MDA-MB-231 and MDA-MB-435 cells were maintained in low-glucose DMEM. All media were supplemented with 10% FBS (Sigma), 1% L-glutamine, 1% of penicillin and 1% streptomycin (GIBCO by Life Technologies, Grand Island, NY). MDA-MB-231 cells with stable transfection with inducible p53-targeting engineered microRNAs were described previously (22) and cultured in low-glucose DMEM with the absence or presence of doxycycline (10 µg/ml) to induce p53 silencing.

The wildtype full-length integrin β4 construct was obtained from Dr. Livio Trusolino (University of Torino, Italy) and described previously (23). The integrin β4 truncated construct, β4 1355T that lacks of signaling domain was amplified by PCR using high fidelity pfu DNA polymerase and cloned into the EcoRI and Sal I sites of pBabe-puro vector. The primers for ITGB4-1355T are: forward (EcoR1), 5’ CAT TAA GAA TTC TAT GGC AGG GCC ACG CCC CA 3’; and reverse (Sal1), 5’ GTA TAT GTC GAC GCG TAG AAC GTC ATC GCT GTA CAT AAG 3’. For stable expression of full-length integrin β4 or β4 1355T in BT549 cells, cells were stably transfected with the empty vector alone or integrin β4 constructs using lipofectamine 2000 and selected with 2 µg/ml of puromycin. The puromycin resistant cells were isolated and the surface expressions of integrin β4 were assessed by fluorescence activated cell sorting using the human integrin β4 antibody (BD Biosciences, clone 439-9B).

For transient gene suppression by siRNA electroporation, cells (3 × 10^6^) from 70% confluent cultures were trypsinized, rinsed with DMEM and electroporated with 200 nM Dharmacon SMARTPool siRNAs specific for an individual target or a control (non-targeting) sequence (Dharmacon, Inc.) as reported previously (24). Cells were cultured normally for 24-96 hrs and then assessed for target gene expression using immunoblotting analysis.

Cisplatin, NU7441, NU7026, KU-55933, and VE-821 were purchased from Selleckchem and doxorubicin from Sigma. For cisplatin treatment, cisplatin at the indicated concentrations was added to cells under normal culture conditions. For inhibitor treatment, cells were pretreated with different inhibitors for 1 hr, then cisplatin was added for additional 24 hr in the presence or absence of inhibitors, as indicated.

### Subcellular protein fractionation and immunoblot analysis

Subcellular fractionation was performed using the Subcellular Protein Fractionation Kit (Thermo Scientific) according to the manufacture’s instruction. For total cell lysates, treated cells were harvested in lysis buffer with phosphatase inhibitors (20 mM Tris-HCl (pH 7.5), 150 mM NaCl, 1 mM Na_2_EDTA, 1 mM EGTA, 1% Triton X-100, 2.5 mM sodium pyrophosphate, 1 mM β-glycerophosphate, 1 mM sodium orthovanadate, 1 µg/ml leupeptin, 1 mM PMSF), sonicated and total cell lysates were subjected to SDS-PAGE, transferred to PVDF membrane and immunoblotted with various antibodies (Cell Signaling Technology). β-actin (monoclonal antibody; Sigma) was used as a loading control for total lysates, and tubulin (Millipore-Sigma) for cytosolic, p84 (GeneTex) for nuclear and histone H2B (Cell Signaling Technology) for chromatin bound fractions.

### UV DNA damage repair analysis

Immuno-slot-blot analysis was performed as described previously (25). Briefly, cells were plated in 60 mm dishes coated with or without 5 µg/ml laminin- 1 in complete growth media. Cells were exposed to 30 J/m^2^ and either harvested immediately or medium replaced. Cells were lysed (10 mM Tris pH 8.0, 1mM EDTA, 0.05% SDS, 100 μg/ml fresh proteinase K) at indicated time points and DNA isolated. DNA was bound to a nitrocellulose membrane using a slot blot apparatus. Membranes were probed using antibody for 6-4PP (Cosmobio) and results presented as percent repair compared to the amount of initial damage (0 hr time point).

### MTT assay

Cells (2 × 10^3^) were seeded in each well of 96-well plate the day before treatments as noted. MTT assays were performed in triplicate or greater by adding 20 µl MTT (5 mg/ml) to each well and incubated at 37°C for 3 hrs. To dissolve the formazan precipitate, 100 µl of stop solution containing 90% isopropanol and 10% DMSO was added and plates agitated for 20 mins at room temperature and then OD 570 was read.

### Immunocytochemistry and the proximity ligation assay (PLA)

BT549 EV and β4 cells (2.5 × 10^4^) were seeded on glass coverslips coated with 5 µg/ml laminin-1 overnight and then treated with 10 µM cisplatin for 24hrs. For immunocytochemistry, cells were then fixed, permeabilized, and immunostained as described previously (26) using the following antibodies: p-p53 S15 and p-53BP1 S1778 (Cell Signaling, Danvers, MA, USA), Cy3- and Cy2-conjugated donkey anti-mouse IgG (Jackson Immune Research, West Grove, PA, USA). DAPI was used to stain nuclei. For PLA assays, cells were fixed and permeabilized according to the Duolink® PLA Fluorescence Protocol (Sigma Aldrich). Primary antibodies used were mouse or rabbit anti-p53 (1:100, Cell Signaling), mouse anti-DNA-PKc (1:100, Cell Signaling), and rabbit anti-53BP1 (1:100, cell signaling). PLA assays were carried out with Duolink® In Situ Detection Reagents Orange (#DUO92007, Sigma Aldrich), Duolink® In Situ PLA® Probe Anti-Rabbit PLUS/MINUS (#DUO92002/DUO92005, Sigma Aldrich) and Duolink® In Situ PLA® Probe Anti-Mouse PLUS/MINUS (#DUO92001/DUO92004, Sigma Aldrich). Cells were imaged using a Nikon Eclipse Ti2 Confocal microscope and Nikon NIS Elements software version 3.2.

### DNA repair reporter assays

BT549 cells (EV and β4, 6 × 10^6^) were electroporated (350V, 500 µF capacity) with 4 µg pDRGFP (HR reporter, Addgene) or 4 µg pimEJ5GFP (NHEJ reporter, Addgene) plus 1.6 µg of pmCherry (transfection control) in the presence or absence of 4 µg pCBASce-I plasmid (Addgene), which expresses I-SceI endonuclease that creates DSB. Upon repair of the reporter, cells express GFP. After treatment with 5 µM cisplatin for 24 hrs, cells (1×10^4^ for each transfection) were analyzed by flow cytometry. The percentage of GFP cell in pmCherry- positive cells was used as the indication of DNA repair efficiency for HR or NHEJ.

### Cell cycle analysis by propidium iodide staining

BT549 cells (EV, β4) were plated on laminin- 1 coated plates and treated with 10 µM cisplatin for 24 hrs. Cells were then trypsinized, rinsed with cold PBS, fixed with cold 70% ethanol, rinsed and then resuspended in PBS staining buffer containing 20 µg/ml propidium iodide I, 0.1% Triton X-100, and 200 µg/ml RNase A and incubated at room temperature for 30 min before analyzed the cell cycle distribution by flow cytometry.

### Statistical analysis and rigor

Data from *in vitro* experiments were compared and analyzed using a two-tailed unpaired Student’s t-test. All experiments were performed at least three times and the representative data are shown. Data were presented as mean ± SD, unless stated otherwise. P values <0.05 between groups were considered significantly different.

## Results

### Integrin α6β4 signaling stimulates NER in response to UV-induced DNA damage

To test the impact of integrin α6β4 on DNA repair in TNBC, BT549 and MDA-MB-435 cells expressing integrin β4 or EV control were seeded onto laminin 1–coated plates, irradiated with UV light, and assessed for DNA repair by resolution of 6-4 photoproducts (6-4PP) as an example of NER. As shown in Figure 1, we found that cells expressing integrin α6β4 rapidly initiated and repaired 6-4PP with a half-life of less than 1 hr while EV control cells took markedly longer. To test whether the integrin α6β4 signaling is involved in the UV-induced DNA damage response, we plated cells without laminin-1 and found no difference in the UV-induced DNA repair kinetics between BT549 EV and β4 cells (**Fig. 1C**), thus demonstrating that the integrin requires its ligand to signal to DNA repair. These data demonstrate that integrin α6β4 signaling promotes NER in TNBC, as we have previously shown in pancreatic cancer (21).

**Figure 1.**
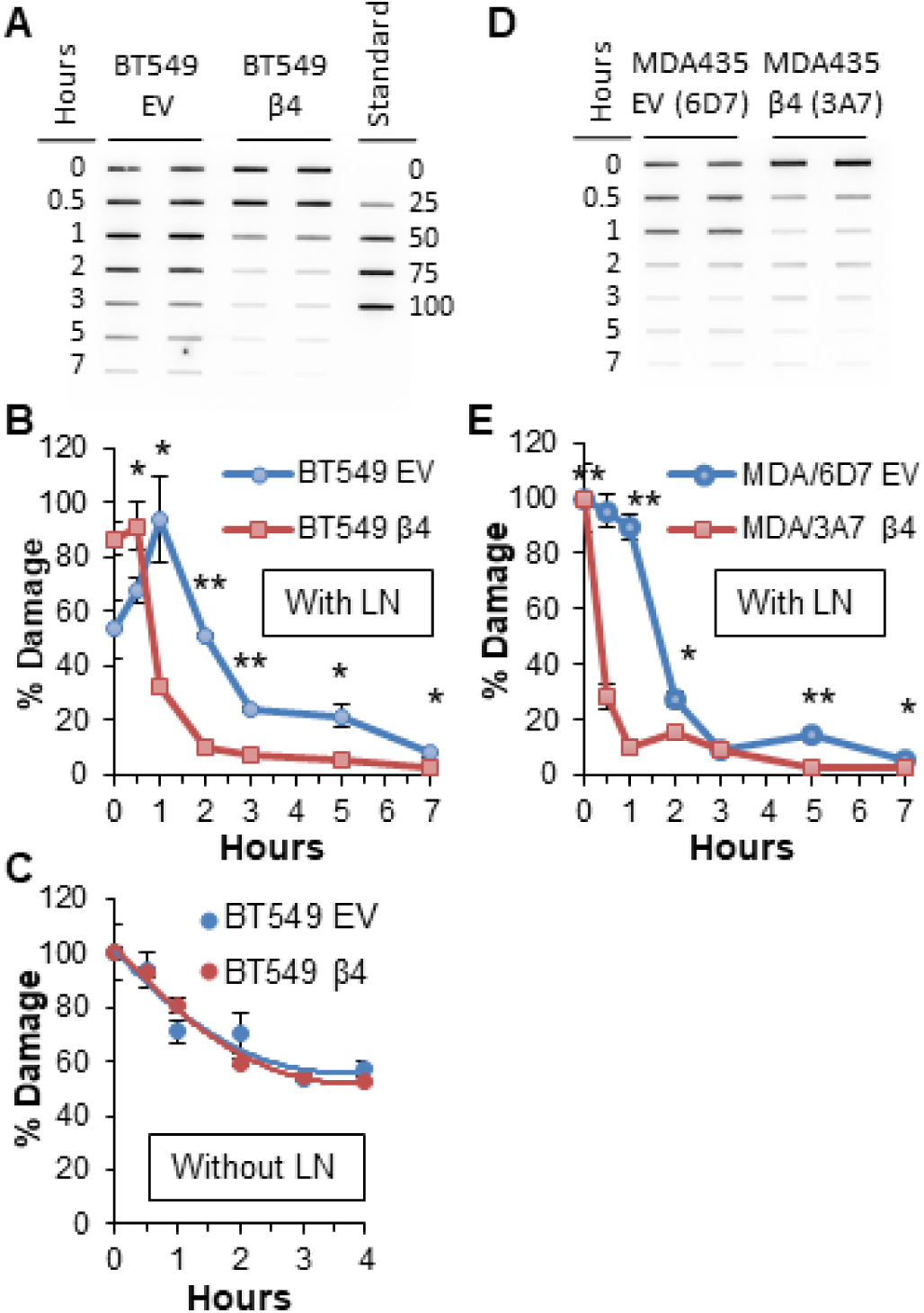
Integrin α6β4 enhances kinetics of nucleotide excision repair (NER) in response to UV irradiation. (A-C) BT549 cells (EV and β4) were plated with (A, B) or without (C) laminin- 1 (LN) and irradiated with 30 J/m^2^ UV light. At the indicated times post irradiation, DNA was extracted, slot blotted and probed for 6-4PP. (D, E). MDA-MB-435 cells (EV-clones 6D7 and β4-3A7) were treated as in (A). (A, D) Representative immunoslot blot of 6-4PP. (B, C, E) Representative immunoslot blot quantification. Data represent 4 separate experiments for each cell line. *p<0.05, **p<0.005.

Next, we sought to identify the DNA damage response (DDR) pathways impacted by integrin α6β4 signaling. DNA damage is sensed through ATR and ATM kinases that in turn phosphorylate a variety of substrates involved in DNA repair including the checkpoint kinases Chk1 (ATR) and Chk2 (ATM). These kinases activate p53 and other downstream effectors to stimulate DNA repair (29,30). To determine how the UV-induced DDR pathway is affected by integrin α6β4 signaling, we irradiated BT549 EV and integrin β4 cells and harvested cells at indicated time points post UV irradiation to assess the phosphorylation and thus activation of key molecules in the DDR pathway. We discovered that integrin α6β4 signaling enhanced phosphorylation of ATM/ATR substrates upon UV irradiation, both in speed and amplitude of response (**Fig. 2A-C**). To identify the critical signaling pathways that are involved in UV-induced integrin α6β4 signaling mediated DNA repair, we immunoblotted these lysates with antibodies against the specific proteins in the DDR pathway. We found that integrin α6β4 signaling dramatically activated ATM, p53, 53BP1 and H2AX (**Fig. 2D**) as evidenced by their enhanced phosphorylation.

**Figure 2.**
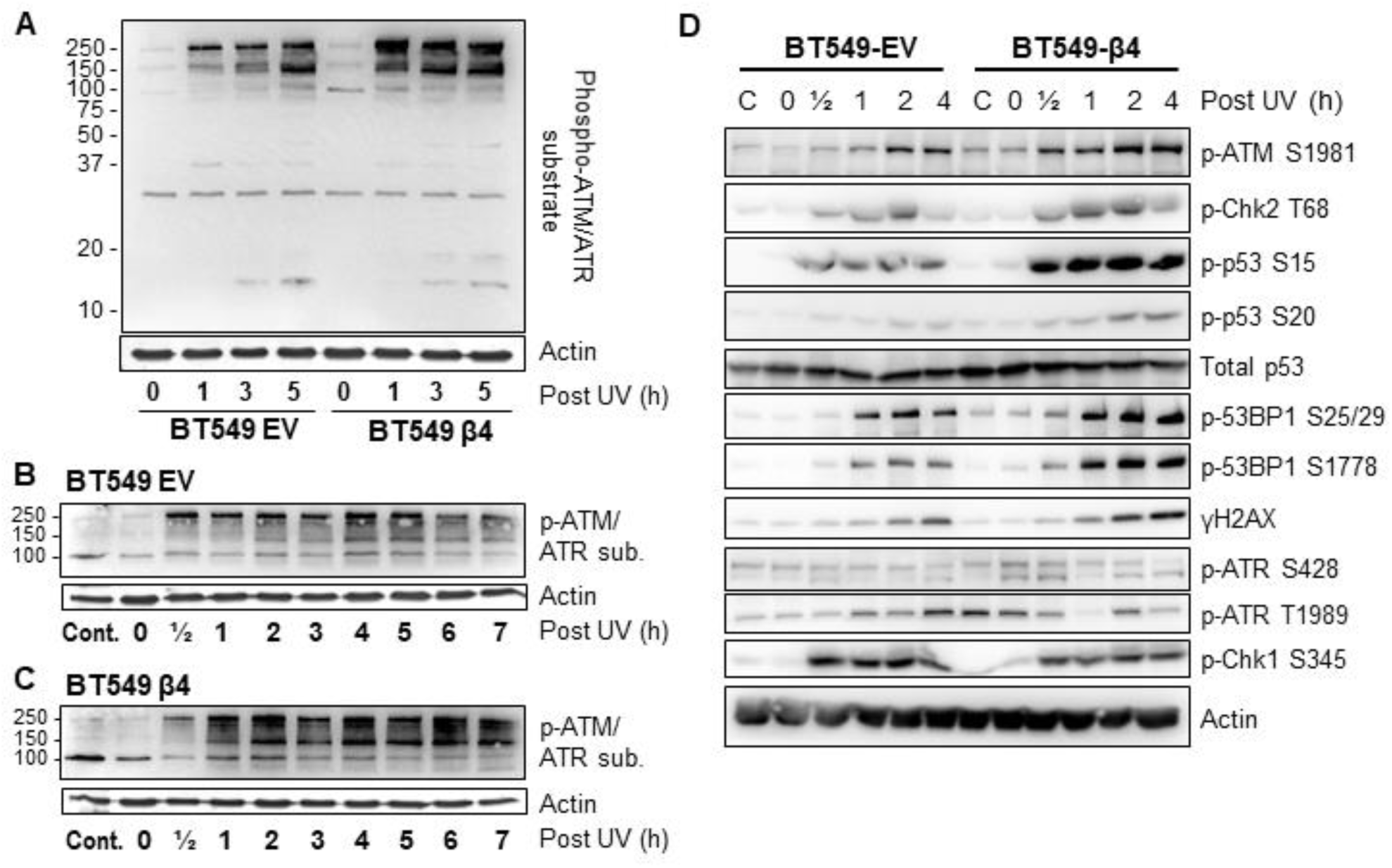
Integrin α6β4 enhances the speed and amplitude of UV-induced DNA damage response. BT549 cells (EV and β4) plated on laminin-1 were UV irradiated and collected for total protein at the indicated time post-UV irradiation. Cell lysates were blotted for phospho- ATM/ATR substrate (A-C) or select proteins in the DNA repair pathways (D). Panel A shows representative time points as a direct comparison while (B) and (C) show full time courses.

### Integrin α6β4 sensitizes TNBC cells to cisplatin treatment

Given that integrin α6β4 signaling impacts DNA repair pathways and most chemotherapies work by generating DNA damage, we hypothesized that integrin α6β4 signaling could influence the response of breast cancer cells to chemotherapy. Accordingly, we treated the BT549 EV and β4 cells with various doses of chemotherapeutic agents including cisplatin, doxorubicin, gemcitabine, and 5-FU for 6 days and then performed MTT assays to assess cell viability. Surprisingly, we found there was a three-fold greater sensitivity to cisplatin in cells expressing the integrin β4 (**Fig. 3A** and **3B**; 1.1 μM LD_50_ for EV vs 0.4 μM for β4). Similar results were obtained in MDA-MB-435 cells (data not shown). However, integrin α6β4 signaling had no effect on the response to gemcitabine (**Fig. 3C**), doxorubicin (**Fig. 3D** and **3E**), and 5-FU (**Fig 3F**).

**Figure 3.**
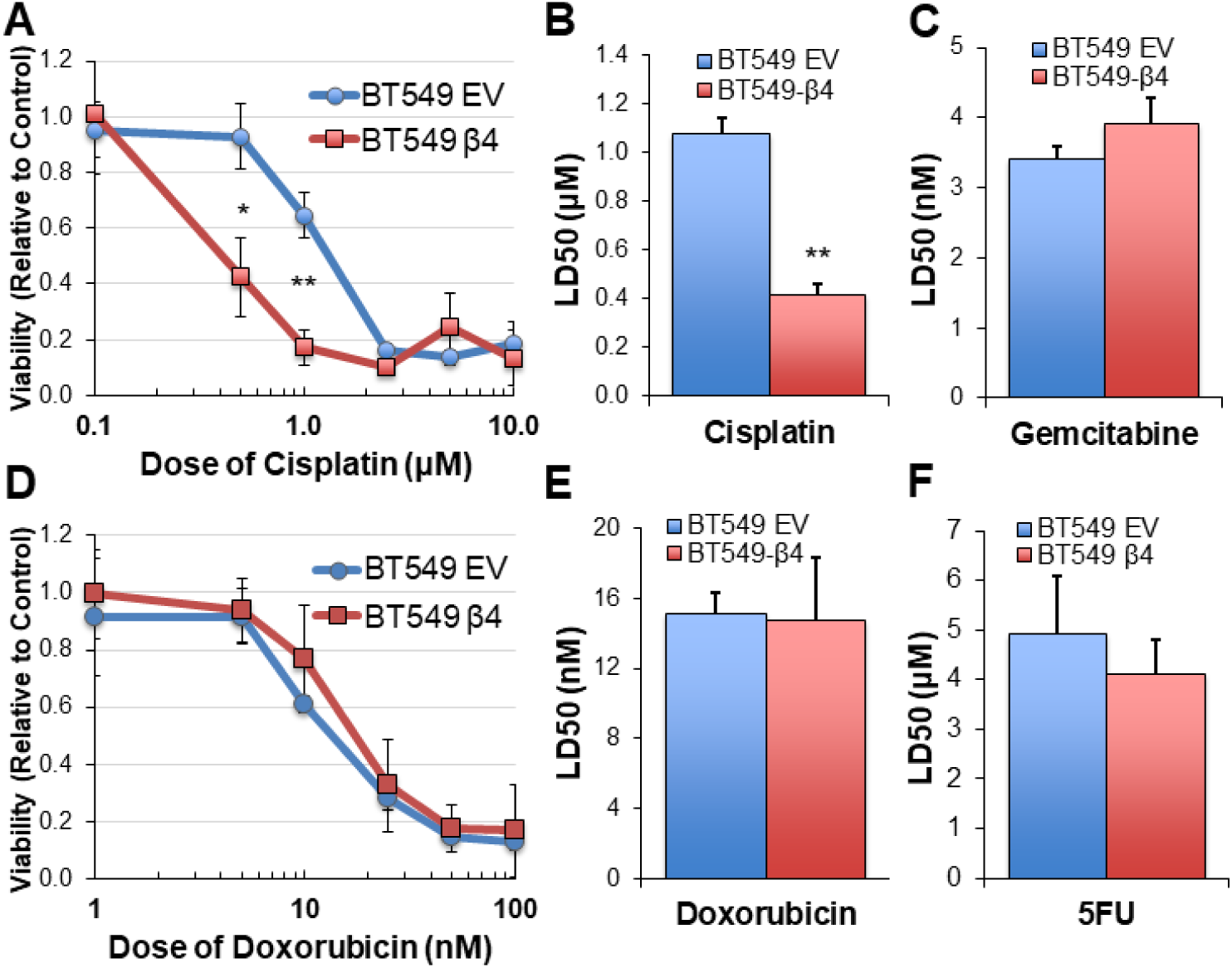
Integrin α6β4 signaling sensitizes BT549 cells to cisplatin treatment. BT549 cells (EV and β4) were treated with various doses of chemotherapeutic agents as indicated for 6 days. Cell viability was assessed by MTT assays and LD50 was calculated for cisplatin (A, B), gemcitabine (C), doxorubicin (D, E) and 5FU (F). *p<0.001, **p<0.0001.

### Integrin α6β4 signaling promotes ATM-p53-53BP1 activation and the association of p53 and 53BP1 with chromatin in response to cisplatin treatment

To determine the impact of integrin α6β4 on cisplatin-mediated DDR signaling, we performed cisplatin dose-response and time-course analyses on BT549 EV and β4 cells. These experiments demonstrated that integrin α6β4 signaling dramatically enhanced the amplitude (**Fig. 4A**) and speed (**Fig. 4B**) of ATM, p53, 53BP1, and H2AX phosphorylation, as well as enhanced PARP1 cleavage, in response to cisplatin treatment. Sensitivity to cisplatin and enhanced PARP1 cleavage as a result of integrin α6β4 seemed unexpected based on its role in promoting cell survival and signaling through the Erk and Akt cell survival pathways (17,31). Furthermore, these pathways can contribute to cisplatin resistance (32,33). Therefore, we investigated the impact of integrin α6β4 signaling on these two survival pathways in conjunction with cisplatin treatment. We found that while the basal activity of Erk was higher in integrin β4 cells compared to EV cells, ERK activation was suppressed upon cisplatin treatment in the BT549 β4 cells but was enhanced in the EV cells. In contrast, the basal phosphorylation of Akt was lower in the BT549 β4 cells and the activation of Akt in response to cisplatin treatment in BT549 EV cells was marginal at low levels of cisplatin but remained unaltered in the BT549 β4 cells (**Fig. 4C**). To test whether the integrin β4 signaling domain is required for the ATM-p53-53BP1 pathway induced by cisplatin, we generated BT549 cells that stably expressed integrin β4 truncation mutation (β4-1355T) in which the signaling domain is deleted (34). Compared to cells expressing wildtype full-length integrin β4, BT549 β4-1355T cells displayed reduced ATM, p53 and 53BP1 activation that were either similar to or less than the BT549 EV cells. These observations suggest that integrin α6β4, through the signaling domain of β4, enhances the DDR to cisplatin through the activation of ATM- p53-53BP1 pathway.

**Figure 4.**
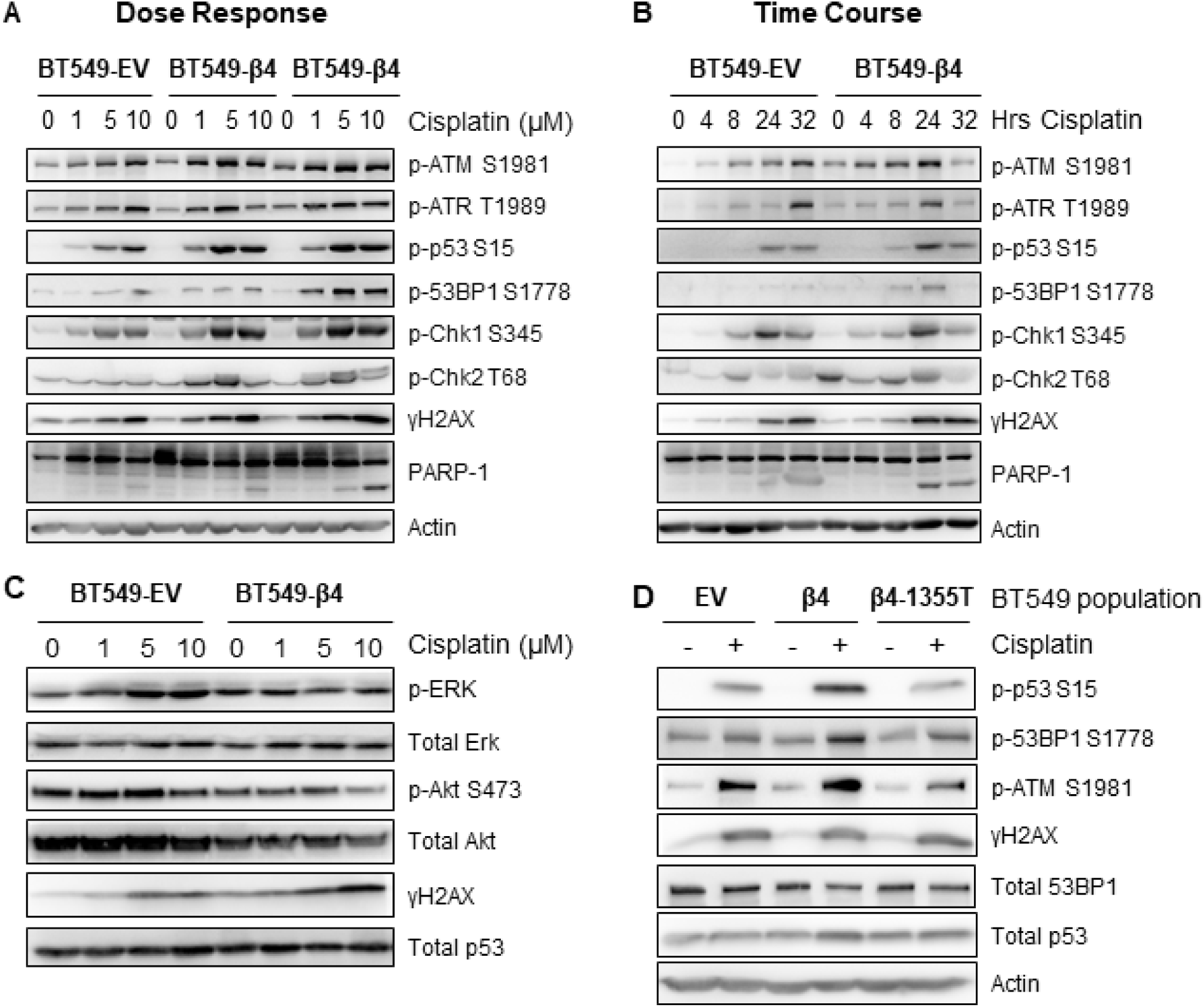
Integrin α6β4 promotes activation of ATM-p53-53BP1 pathway in a time and dose dependent manner in response to cisplatin that results in enhanced DNA damage and PARP1 cleavage. (A) BT549 cells (EV and β4) were plated on laminin-1, treated with the indicated dose of cisplatin for 24 hrs (A) or 10 μM cisplatin for indicated times (B) and assessed for phosphorylation of indicated DDR proteins as noted. (C) Cells treated as in (A) were assessed for phospho-Erk, phospho-AKT, γH2AX, and total Erk, Akt and p53. (D) BT549 cells (EV, β4, and β4-1355T) were plated on laminin-1 and treated with 10 μM cisplatin for 24 hrs prior to immunoblotting with indicated DDR proteins.

Next, we sought to compare how nuclear activation of p53 by cisplatin treatment and integrin α6β4 compared to cytosolic levels using subcellular fractionation. We found that p53 activation in the nucleus was more dramatic than that present in the cytosol (**Fig. 5A**). This enhanced activation of p53, and that of 53BP1, were confirmed by immunocytochemistry (**Fig. 5B**). To further test whether these activated proteins were in the nucleoplasm (soluble) or associated with chromatin, we performed subcellular protein fractionation and immunoblotting for phosphorylated and total p53 and 53BP1. We found that, compared to the EV cells, the associations of both p53 S15 and 53BP1 S1778 with chromatin, as well as γH2AX, were enhanced in the integrin β4 cells in response to cisplatin treatment. Furthermore, soluble γH2AX was dramatically increased in BT549 integrin β4 cells upon cisplatin treatment. Interestingly, total p53 levels associated with the chromatin were amplified with integrin α6β4 regardless of the treatment condition.

**Figure 5.**
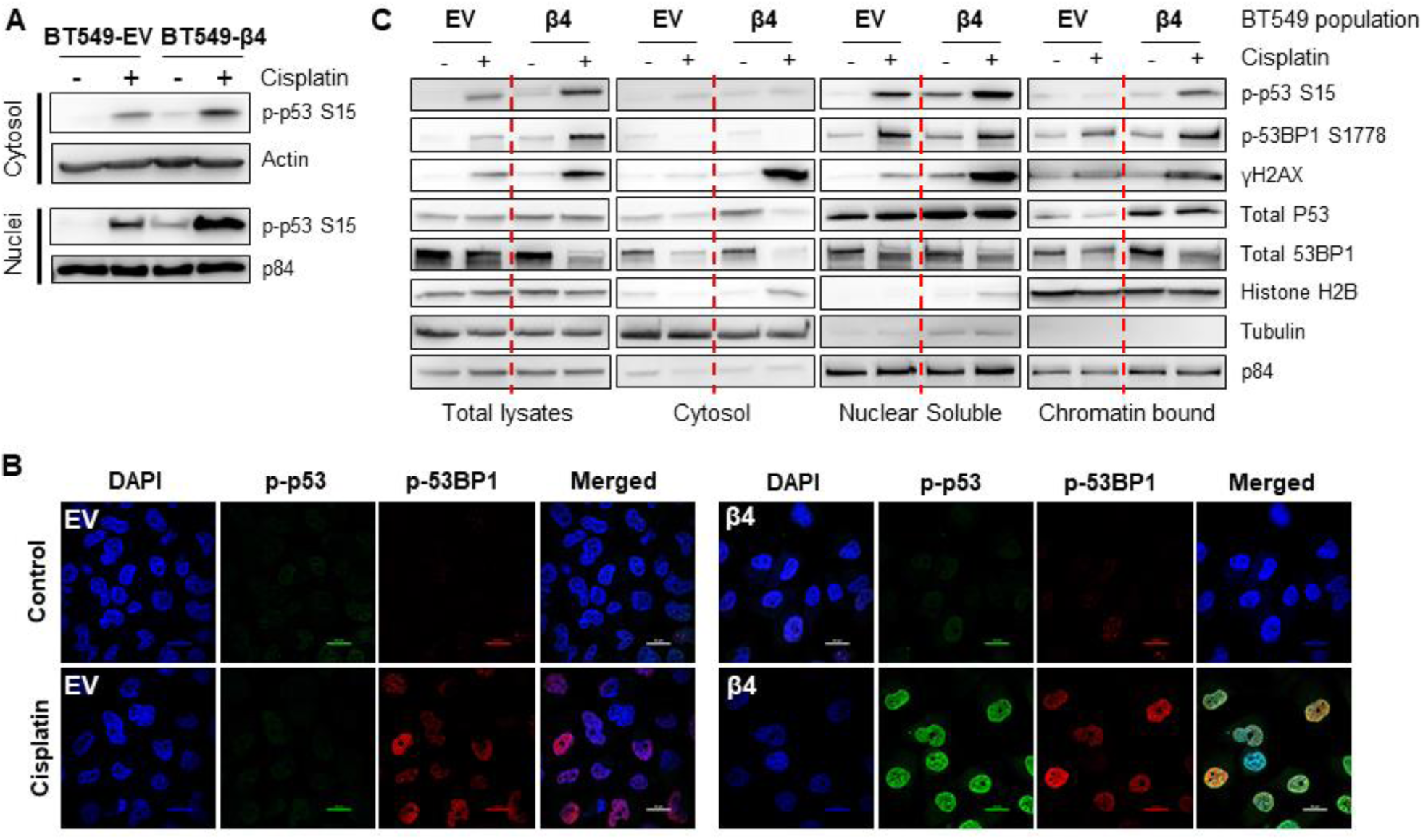
Integrin α6β4 signaling promotes recruitment of p53 and 53BP1 to chromatin in response to cisplatin treatment. BT549 cells (EV and β4) plated on laminin-1 and treated with 10 μM cisplatin for 24 hrs were harvested for cytosolic or nuclear fractions and immuno-blotted for phospho-p53 using actin and p84 as controls (A) or for immunocytochemistry staining for phospho-p53 S15 and phospho-53BP1 S1778 as indicated (B, scale bars, 20µm). (C) Subcellular protein fractionation was performed on cells treated as in (A) and noted fractions were immunoblotted with DDR proteins as indicated. Tubulin was used as the marker for total and cytosolic protein, Histone H2B and p84 were used as the markers for nuclear fractions.

### Mutant p53 is required for integrin α6β4-mediated ATM/53BP1/p53 pathway activation and cisplatin sensitivity

Mutation rates for p53 are high in TNBC (85% (35)) where they tend to co-exist with high integrin α6β4 expression (cBioPortal analysis, p<0.001). Therefore, we sought to test whether integrin α6β4 cooperates with mutant p53 to alter cisplatin sensitivity. Thus, we attained MDA- MB-231 cells with doxycycline-inducible knockdown of mutant p53 (36), induced suppression of p53 with doxycycline and/or knocked down integrin β4 expression by siRNA or left cells untreated during cisplatin treatment. The data revealed that, compared to the knockdown of integrin α6β4 or p53 alone, the effect of knockdown of both mutant p53 and integrin β4 (**Fig. 6B**) on cell viability is additive (**Fig. 6A;** p = 0.03) and highly significant (p < 0.0001 vs control at 1 μM and 2.5 μM). To test the requirement for mutant p53 in cisplatin-induced DDR, we knocked down p53 by siRNA in BT549 EV and integrin β4 cells, treated these cells with cisplatin and then assessed DNA repair pathways. We show that knockdown of p53 blocked activation of ATM in response to cisplatin and/or integrin α6β4 as well as the downstream 53BP1 phosphorylation (**Fig. 6C**). Collectively, these results demonstrate that mutant p53 is required for the amplification of ATM activity and 53BP1 phosphorylation downstream of integrin α6β4 signaling in response to cisplatin.

**Figure 6.**
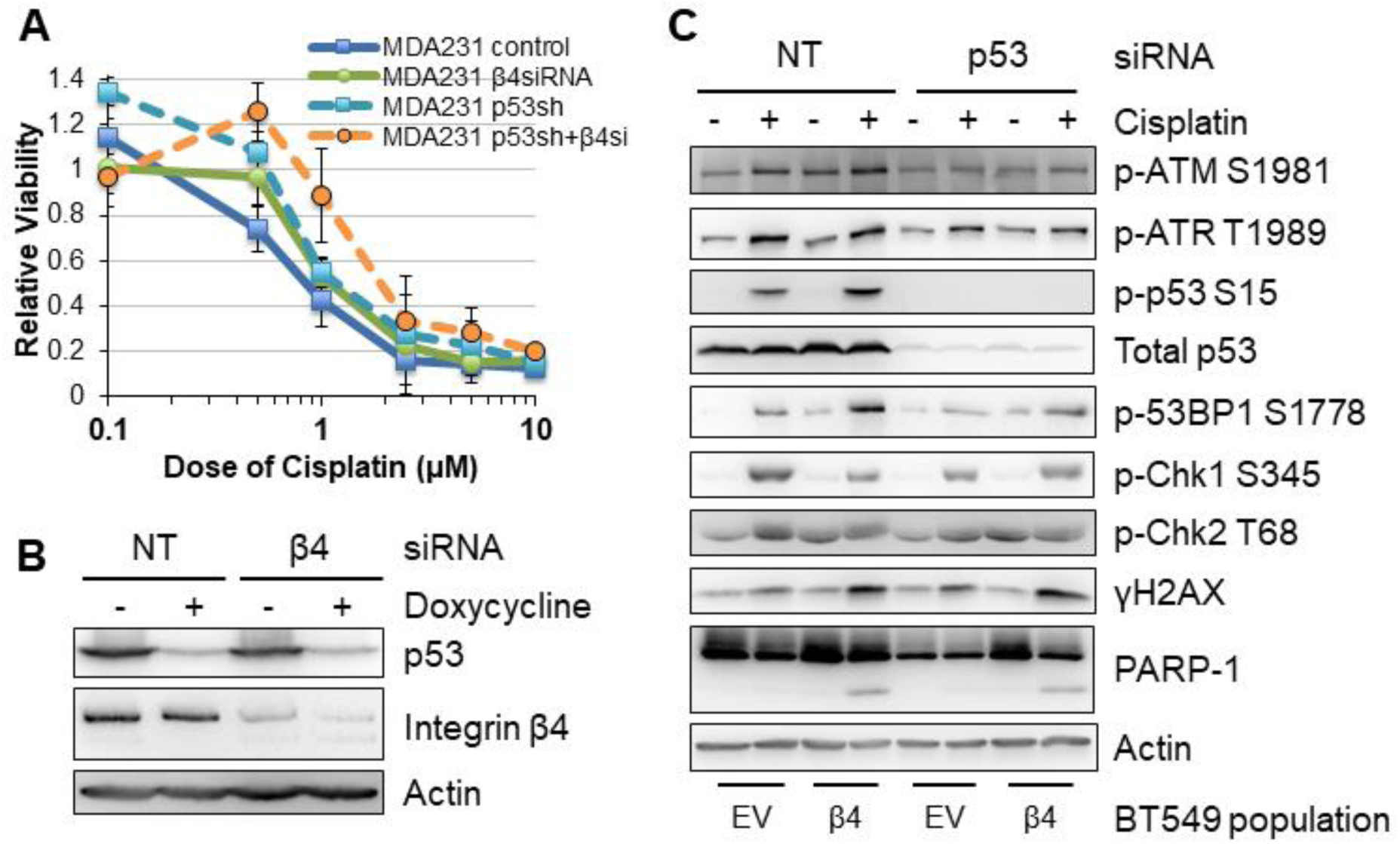
Mutant p53 is required for integrin α6β4-mediated sensitivity to cisplatin and activation of ATM and 53BP1. (A and B) Mutant p53 was knocked down in a doxycycline-inducible manner in MDA-MB-231 cells and/or integrin β4 was knocked down by siRNA. Then cells were plated on laminin-1-coated plates, treated with various doses of cisplatin for 6 days. Cell viability was assessed by MTT (A) and the efficiency of knockdown was tested by immunoblotting (B). (C) p53 was knocked down by siRNA in BT549 EV and BT549 β4 cells, then cells were plated on laminin-coated plates and treated with 10 µM cisplatin for 24 hrs prior to harvesting for immunoblotting analysis.

### Inhibition of ATM and/or ATR differentially affects UV and cisplatin induced DDR signaling downstream of integrin α6β4

To test the impact of ATM and ATR kinases on DDR signaling downstream of integrin α6β4 signaling after UV damage and cisplatin treatment, we plated BT549 cells (EV and β4) on laminin-1 and pretreated cells with inhibitors for ATM (KU55933), ATR (VE-821), both ATM and ATR (M/R) or carrier for 1 hr. Cells were then UV irradiated and incubated for 3 hrs or treated with cisplatin for 24 hrs. Cell lysates were immunoblotted to assess the activation of key proteins in the DDR pathway. We found that with UV irradiation, integrin α6β4 signaling substantially enhanced p53 (S15) and 53BP1 (S25/29, S1778) phosphorylation downstream of both ATM and ATR, as it required both inhibitors to suppress these events. Integrin α6β4 also enhanced phosphorylation of Chk1, Chk2 and p53 (S20) (**Fig. 7A**). As expected, Chk1 was most sensitive to ATR inhibition while Chk2 was inhibited by ATM inhibition. Interestingly, ATM inhibition blocked p53 (S20) phosphorylation but not ATR, suggesting this signaling predominates through the ATM- Chk2 pathway. Notably, these phosphorylation events in response to cisplatin treatment were similar between BT549 EV and β4 cells with the exception of 53BP1 phosphorylation, which remained high in the presence of ATM and ATR inhibitors (**Fig. 7B)**.

**Figure 7.**
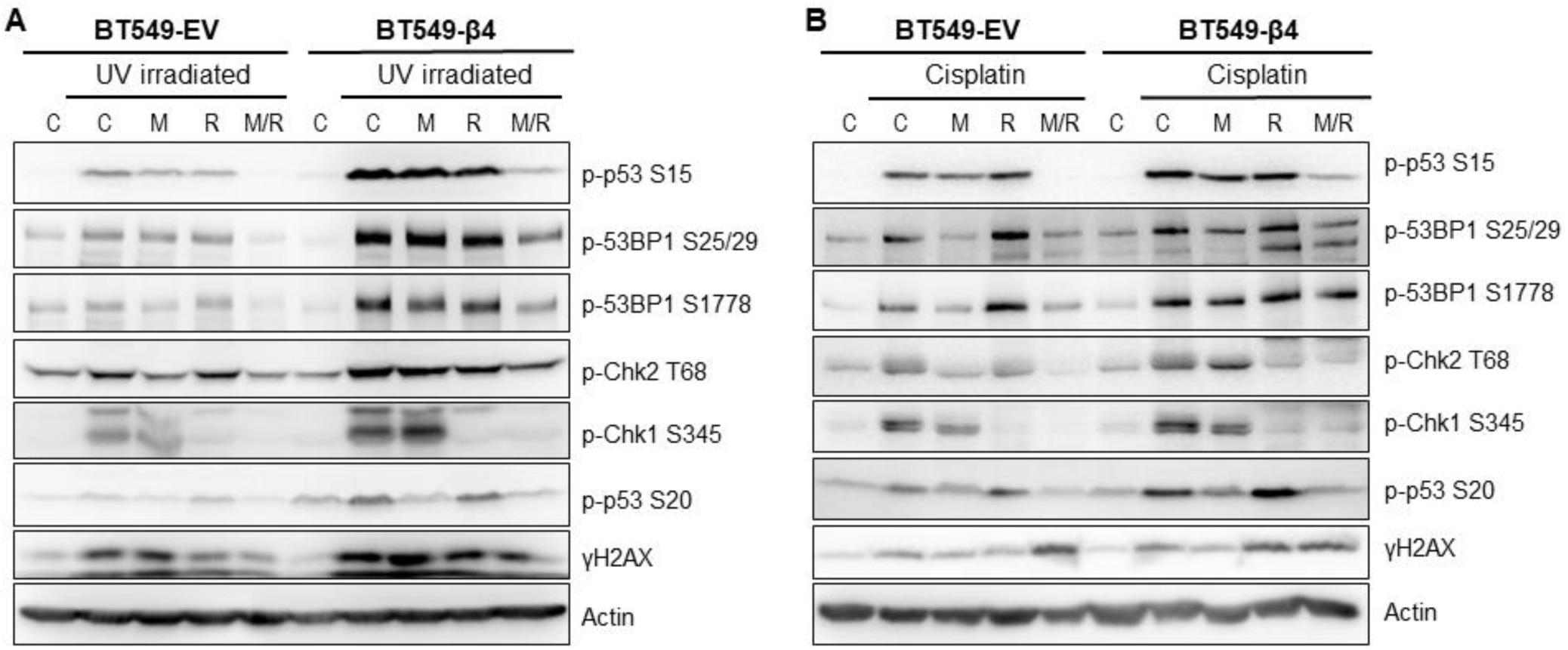
Sensitivity of DNA damage response signaling to ATM or ATR inhibition depends on source of damage and integrin α6β4 status. BT549 cells (EV and β4) were plated on laminin-1-coated plates, pretreated with 10 µM KU-55933 (ATM inhibitor, M), VE-821 (ATR inhibitor, R) or the combination of both inhibitors (M/R) for 1 hr before UV irradiated (A) or 10 µM cisplatin treatment (B). Cell lysates from 3 hrs post UV irradiation or 24 hrs post cisplatin treatment were immunoblotted with signaling proteins in DNA-repair pathway as noted.

### DNA-PKc is preferentially activated downstream of integrin α6β4 and required for enhanced 53BP1 phosphorylation

Chk2 (37), p53, and 53BP1 (38) are targets of DNA-PKc, a DNA damage sensing kinase involved in NHEJ DSB repair. To test whether DNA-PKc is involved in integrin α6β4 enhanced DDR, we assessed how cisplatin and integrin α6β4 signaling impact DNA-PKc activation and the influence of DNA-PKc inhibition on downstream DDR signaling. As shown in Figure 8, cisplatin treatment resulted in DNA-PKc phosphorylation at S2056 and T2609, which are indicative of an activated kinase; this activation was substantially greater in BT549 β4 cells than in EV cells. To determine how DNA-PKc activity affects cisplatin-induced DNA repair pathways, we pretreated BT549 EV and integrin β4 cells with DNA-PKc inhibitors NU7441 or NU7026 at various concentrations prior to cisplatin treatment. We find that phosphorylation of 53BP1 in response to cisplatin treatment was particularly sensitive to DNA-PKc inhibition, suggesting that DNA-PKc controls 53BP1 phosphorylation. We noted that at higher concentrations of inhibitor, ATM and ATR activities were impacted as well as the checkpoint kinases and p53, suggesting these molecules were impacted by DNA-PKc indirectly or were inhibited non-specifically at these drug concentrations. We next investigated DNA-PKc-p53, p53-53BP1, and DNA-PKc-53BP1 complexes by PLA with and without cisplatin treatment. We found that DNA-PKc-p53 complexes and p53-53BP1 complexes formed preferentially in the integrin β4 expressing cells after cisplatin treatment; however, DNA-PKc did not appear to complex directly with 53BP1. These data, coupled with our observation that mutant p53 was required for 53BP1 activation (**Fig. 6**), suggest that integrin α6β4 signaling to DNA-PKc controls 53BP1 phosphorylation in response to cisplatin by activating and recruiting p53 to link DNA-PKc to 53BP1.

**Figure 8.**
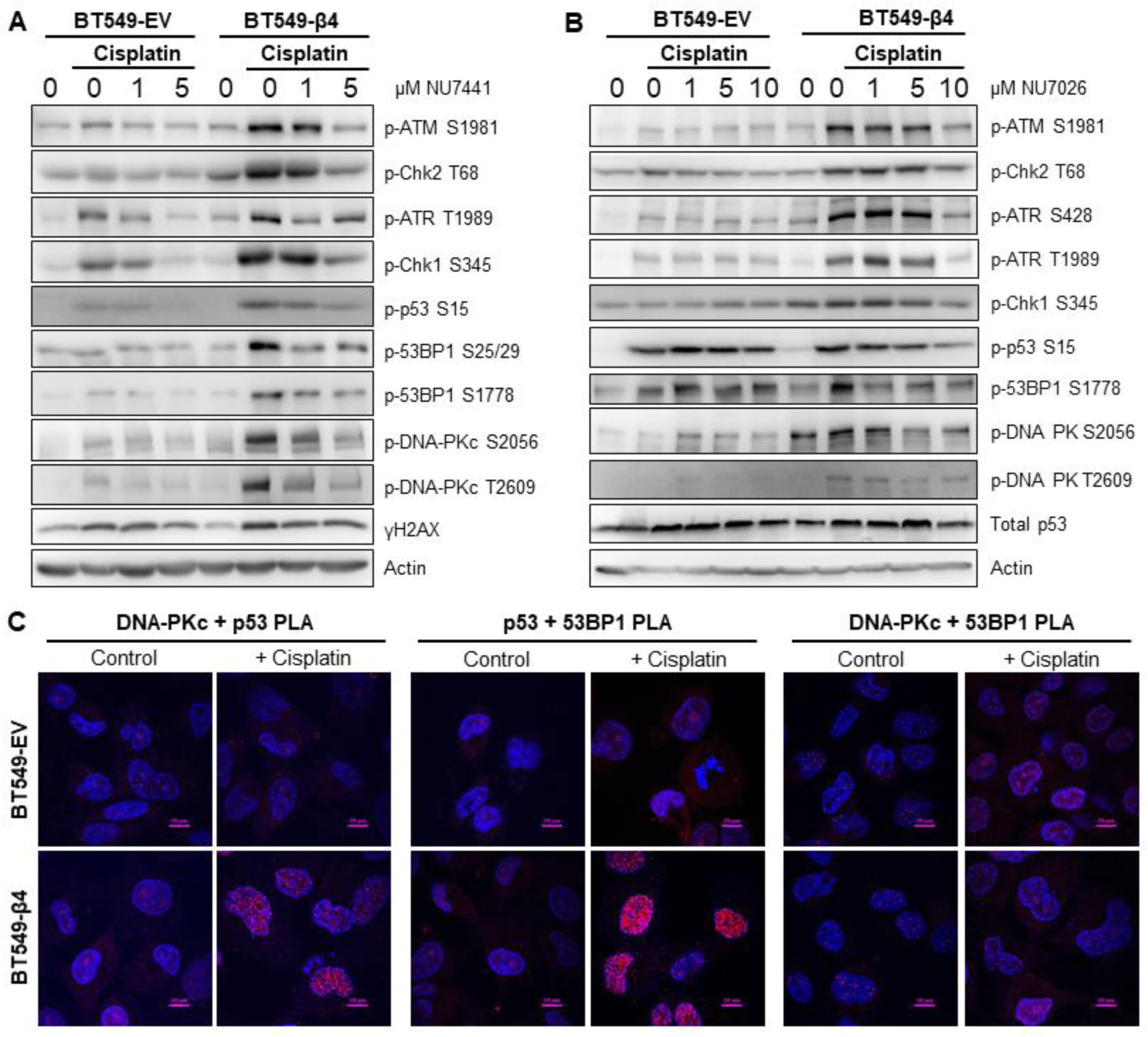
Effect of DNA-PKc inhibition on cisplatin-induced DNA repair pathway. BT549 cells (EV and β4) were plated on laminin-1-coated plates, pretreated with DNA-PKc inhibitors NU7441(A) or NU7026 (B) at indicated concentrations for 1 hr before treatment with 10 µM cisplatin for 24hrs. Cell lysates were then immunoblotted for signaling proteins in DNA repair pathway as noted. (C) BT549 cells (EV and β4) plated on laminin-1-coated coverslips were treated with 10 µM cisplatin for 24hrs, the associations of DNA-PKc, p53, and 53BP1 were assessed by PLA, as noted. Scale bars, 10µm.

### Integrin α6β4 switches DSB repair from HR to NHEJ

DNA-PKc is the DNA damaging sensing kinase that is involved in the NHEJ DSB repair. Furthermore, 53BP1 is important for the switch between HR and NHEJ that can result in enhanced cisplatin sensitivity (39). Collectively, our data suggests that integrin α6β4 signaling could shift DSB repair from HR to NHEJ. To test this hypothesis, we utilized HR and NHEJ reporter systems that use the endonuclease Sce-1 to cause DSBs that, upon repair, create a functional GFP molecule. Here, BT549 EV and integrin β4 cells were co-transfected with pDR GFP (HR reporter) or pimEJ5GFP (NHEJ reporter) in the presence or absence of pCBASce-I and with pmCherry as an internal transfection control. Transfected cells were then treated with or without 5 μM cisplatin for 48 hrs and analyzed for GFP and mCherry by flow cytometry. As shown in Figure 9, BT549 EV cells had a higher basal HR activity than the integrin β4 cells. Conversely, the integrin β4 cells displayed higher basal level of NHEJ activity than the EV cells. Upon cisplatin treatment the BT549 EV cells dramatically activated HR activity but suppressed HR in integrin β4 cells. In contrast, NHEJ activity in integrin β4 cells was dramatically activated upon cisplatin treatment compared to the EV cells. Since NHEJ is known to function as the primary DSB repair mechanism in G1, we investigated the cell cycle distribution of cells under these conditions. We found that in untreated cells that the cell cycle distribution was similar between the EV and β4 expressing cells. When the cells were treated with cisplatin, both cell populations showed a 2.5-fold increase in S phase and a concomitant drop in G1 distribution. In contrast, BT549 EV cells lost approximately 40% of their G2 distribution, while the β4 cells doubled their G2 distribution. These data suggest that cell cycle distribution, such as a blockade in G1, does not play into the switch between HR and NHEJ pathways downstream of integrin α6β4.

**Figure 9.**
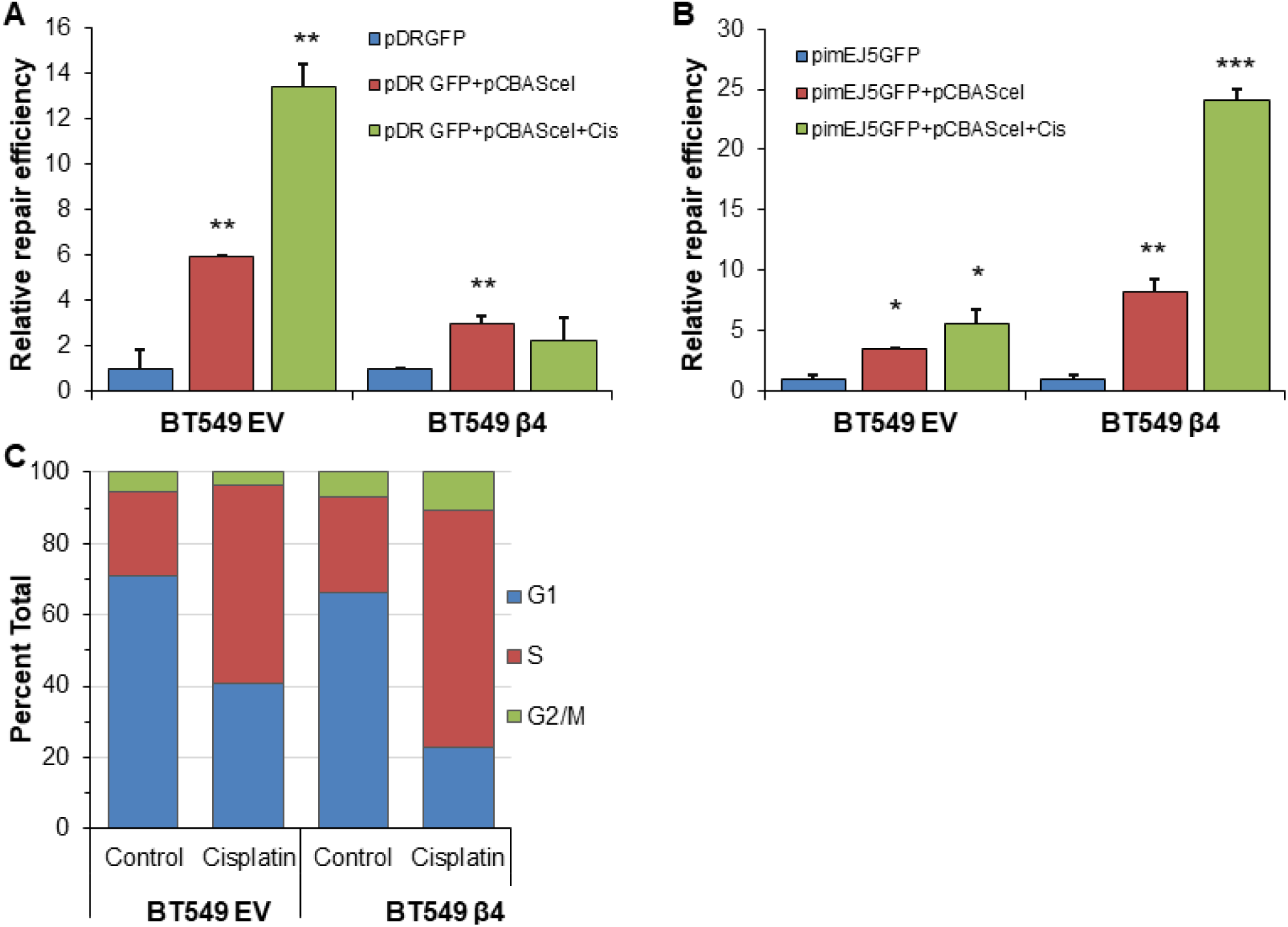
Integrin α6β4 signaling enhances NHEJ and suppresses HR. BT549 cells (EV, β4) were electroporated with pDRGFP (HR reporter; A) or pimEJ5GFP (NHEJ reporter; B) in the presence or absence of pCBASce-I plasmid, which expresses I-SceI endonuclease that causes DSB and plated on laminin-1 coated plates. Upon repair of the reporter, cells express GFP. After treatment with 5 µM cisplatin for 24 hrs, cells (1×10^4^ for each transfection) were analyzed by flow cytometry using cotransfected pmCherry as a transfection control. (C) BT549 cells (EV, β4) plated on laminin-1 coated plates were treated with 10µM cisplatin for 24 hrs and assessed for cell cycle distribution using propidium iodide staining and flow cytometry analysis. *p<0.05, **p<0.001, ***p<0.0001.

## Discussion

In this study, we demonstrate that integrin α6β4 enhances DDR signaling (**Figs. 2,4,7-8**), stimulates DNA repair (**Fig. 1**), shifts DSB repair from HR to NHEJ (**Fig. 9**), and promotes cisplatin sensitivity (**Fig. 3**) as a result of signaling through DNA-PKc (**Fig. 9D**). These processes are dependent on the ability of integrin α6β4 to activate and signal through mutant p53 (**Fig. 6**). These observations are interesting considering it is wild-type p53 that is associated with apoptosis in response to DNA damage. Notably, the clinical significance of p53 depends on its mutational status where a mutant p53 gain-of-function gene signature correlates best with clinical outcomes (40). We find in the literature (41,42) and our data (**Figs. 3, 6**) that cisplatin sensitivity is higher in a mutant p53 background when integrin α6β4 is expressed, thus suggesting that integrin α6β4 can control mutant p53 response to platinum-based therapy.

Cisplatin sensitivity results from three major mechanisms. Platinum agents diffuse through the membrane but are also influxed through copper transporters or effluxed by ABC7 family transporters. Once in the cell, glutathione can sequester platinum in the cytosol and prevent nuclear import. Finally, DNA repair mechanisms can define the final sensitivity. Of these mechanisms, DNA repair produces the lowest fold change in resistance or sensitivity (43). In TNBC, however, HR-mediated DNA repair deficiency has the strongest association with efficacy of platinum-based therapies and is a major determinant of which patients receive platinum regimens (43,44).

HR-deficiency is loosely defined as having mutations or loss of expression (e.g. through promoter methylation) of key molecules within the HR pathway (44). Despite its ability to shuttle DSB repair from HR to NHEJ, DNA-PKc is not generally associated with HR-deficiency conceptually. Our data demonstrate that integrin α6β4 signaling through DNA-PKc contributes to cisplatin sensitivity by promoting phosphorylation of 53BP1 and a shift from HR to NHEJ. Clinically, DNA-PKc inhibitors are ineffective as a monotherapy, but have been found to cooperate with doxorubicin and ionizing radiation (45,46), which are indiscriminate in which DSB repair pathway used. Importantly, we find that inhibition of DNA-PKc with two different chemical inhibitors reverses the sensitivity attributed to integrin α6β4. Given the importance of integrin α6β4 to basal-like breast cancer, these observations give additional rationale that DNA-PKc should be further investigated as a mediator of HR deficiency and cisplatin sensitivity in TNBC.

In breast cancer management, there is considerable interest in expanding what is considered HR-deficiency (44). A recent clinical trial investigating this concept demonstrated that patients with germline mutations in *BRCA1* or *BRCA2* benefited from carboplatin, which creates the same DNA lesions as cisplatin but with different toxicities (47), when compared to docetaxel treatment. This benefit was not observed with *BRCA1* promoter methylation, low BRCA1 mRNA or HR-deficiency defined by the Myriad assay (48). However, when HR-deficiency is defined by the types of DNA damage, cisplatin as a neoadjuvant therapy gave a therapeutic advantage over other chemotherapies (49). Notably, the loss of heterozygosity, telomeric allelic imbalance, and large-scale state transitions used to define HR-deficiency in this study are also types of damage known to be created by the switch from HR to error prone NHEJ (50).

53BP1 is a major down-stream mediator of p53 and DNA-PKc that has been implicated in the decision between HR and NHEJ that can alter sensitivity to cisplatin (39,51,52). Notably, loss of 53BP1 has been attributed to cisplatin resistance in BRCA1 mutant cells (53). While it has been unclear how mutant p53 impacts 53BP1 function (51), our data suggest that mutant p53 brings 53BP1 in close proximity with DNA-PKc to allow it to be phosphorylated on multiple sites to potentiate its function in switching repair to NHEJ. Mutant p53, which is found in 85% of TNBCs (35), has been documented to promote either sensitivity or resistance to cisplatin depending on biological context (40,42,54), including direct blockade of the HR pathway (55). Our data indicate that integrin α6β4 may be instrumental in providing that context and promoting cisplatin sensitivity by suppressing the HR repair and forcing DSB repair to the NHEJ pathway through the mutant p53-53BP1 interactions. The co-occurrence of high rates of mutant p53 and high expression levels of integrin α6β4 in TNBC and basal-like breast cancers could provide rationale for why cisplatin sensitivity predominates in these breast cancer subtypes. These observations collectively suggest that integrin α6β4 expression, DNA-PKc activity, *TP53* mutational status and 53BP1 expression should be considered when defining HR-deficiency and may be important in identifying those patients that will benefit most by neoadjuvant cisplatin as a first line chemotherapeutic.

The involvement of integrin α6β4 in promoting DNA repair and TNBC progression appears to contradict its role in chemosensitivity. However, chemotherapy regimens have evolved to work on the most aggressive and highly proliferative cancers, as exampled by the observation that TNBCs receive clinical benefit from chemotherapy while Luminal A breast cancers do not (56,57). Thus, it is logical that drivers of progression can provide intrinsic sensitivity to specific chemotherapies. Sensitivity to cisplatin and enhanced PARP1 cleavage as a result of integrin α6β4 signaling also seems unexpected based on the integrin’s role in promoting cell survival. Interestingly, Akt and Mek-Erk signaling can be shut down during the DNA damage response while sustained signaling results in resistance (32). We find that Erk is activated in BT549 EV cells with cisplatin treatment. In contrast, Akt and Erk are suppressed in BT549 β4 cells with cisplatin treatment (**Fig. 4C**), thus suggesting integrin α6β4 cannot enhance Akt or Erk signaling to promote survival in response to cisplatin. How integrin α6β4 suppresses these activities is not currently known, but may provide rationale as to why it does not promote resistance to chemotherapies, as previously suggested (20).

Conventional thought is that p53 is downstream of ATM/ATR. However, we find that p53 lies upstream of ATM given our results that p53 is required for integrin α6β4 to amplify ATM phosphorylation. Specifically, our data show that knockdown of p53 prevents activation of ATM beyond a basal level and eliminates the ability of integrin α6β4 to promote ATM activation in response to cisplatin. These data do not imply that ATM is not active, but rather that p53 is responsible for a positive feedback that further amplifies ATM activity, as shown in our summary model (**Fig. 10**). Integrin α6β4 signaling in response to cisplatin treatment also activates DNA- PKc that, in its association with mutant p53, leads to the phosphorylation of 53BP1 and impacts the DNA repair pathway choice. Our data also imply that cell cycle phase does not contribute to the switch from HR to NHEJ downstream of integrin α6β4 signaling in response to cisplatin, but rather the integrin directly signals this switch through the DNA-PKc-p53-53BP1 pathway.

**Figure 10.**
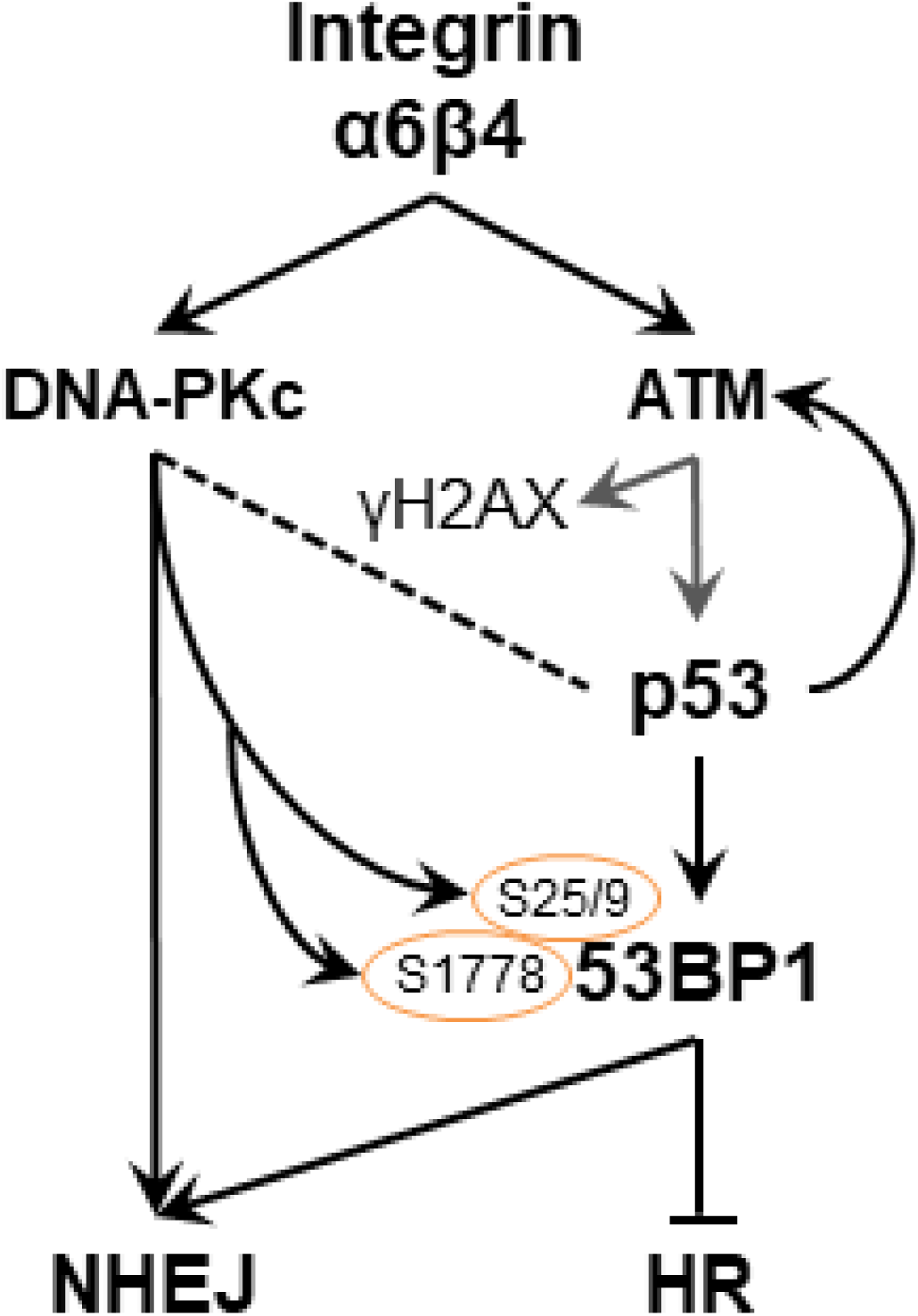
Summary Model.

In summary, we trace the ability of integrin α6β4 to enhance DNA repair and affect cisplatin sensitivity to the ability of integrin α6β4 to signal through mutant p53, amplify ATM and DNA-PKc activity, increase 53BP1 phosphorylation, and switch double strand break repair from HR to NHEJ. This signaling, against the backdrop of prominent losses and gains of select DNA repair molecules in breast cancer as a whole, may lead to a novel HR-deficiency that characterizes TNBC and their response to select chemotherapies.

## Funding

This work was supported by the National Institutes of Health through National Cancer Institute [grant numbers R01 CA223164-01 to KLO, R21 CA178753 to KLO, and R01 CA131075 to JAD], and National Center for Advancing Translational Sciences (grant number KL2 TR001996 to RLS); and by a Markey Women Strong Award through the Markey Cancer Foundation [no grant number, to KLO] and the University of Kentucky Center for Cancer and Metabolism (grant number P20GM121327) for providing services through the Imaging Core and for the pilot funding (no grant number, to MC). The Markey Cancer Center Biospecimen Procurement and Translational Pathology, the Biostatistics and Bioinformatics, and Flow Cytometry and Immune Monitoring Shared Resource Facilities, which supplied services for this study, are supported by National Institutes of Health [grant number P30 CA177558]. The content is solely the responsibility of the authors and does not necessarily represent the official views of the NIH.

## Acknowledgments

We gratefully acknowledge Drs. Eva Goellner, Tadahide Izumi, and David Orren for thoughtful suggestions and discussions regarding this study. We also thank Dr. Jill Bargonetti from Hunter College of the City University of New York and Dr. Livio Trusolino for reagents.

## Conflict of Interest

The authors reported no conflict of interest.

